# Selective retinal ganglion cell loss and optic neuropathy in a humanized mouse model of familial dysautonomia

**DOI:** 10.1101/2021.06.04.447086

**Authors:** Anil Chekuri, Emily M. Logan, Aram J. Krauson, Monica Salani, Sophie Ackerman, Emily G. Kirchner, Jessica M. Bolduc, Xia Wang, Paula Dietrich, Ioannis Dragatsis, Luk H. Vandenberghe, Susan A. Slaugenhaupt, Elisabetta Morini

**Author notes:** Correspondence should be addressed to S.A.S. and to E.M. These authors contributed equally to this work (Susan A. Slaugenhaupt, Elisabetta Morini).

## Abstract

Familial dysautonomia (FD) is an autosomal recessive neurodegenerative disease caused by a splicing mutation in the gene encoding Elongator complex protein 1 (*ELP1*, also known as *IKBKAP*). This mutation results in tissue-specific skipping of exon 20 with a corresponding reduction of ELP1 protein, predominantly in the central and peripheral nervous system. Although FD patients have a complex neurological phenotype caused by continuous depletion of sensory and autonomic neurons, progressive visual decline leading to blindness is one of the most problematic aspect of the disease, as it severely affects their quality of life. To better understand the disease mechanism as well as to test the *in vivo* efficacy of targeted therapies for FD, we have recently generated a novel phenotypic mouse model, *TgFD9; Elp1*^∆20/flox^. This mouse exhibits most of the clinical features of the disease and accurately recapitulates the tissue-specific splicing defect observed in FD patients. Driven by the dire need to develop therapies targeting retinal degeneration in FD, herein, we comprehensively characterized the progression of the retinal phenotype in this mouse, and we demonstrated that it is possible to correct *ELP1* splicing defect in the retina using the splicing modulator compound (SMC) BPN-15477.

## Introduction

Familial dysautonomia (FD, MIM223900), also known as Riley–Day syndrome or hereditary sensory and autonomic neuropathy type III (HSAN III), is a rare, recessive, early onset, sensory and autonomic neurodegenerative disorder caused by T to C nucleotide change in the 5’splice site of intron 20 of the *Elongator complex protein 1* (*ELP1* also known as *IKBKAP*). This mutation leads to tissue-specific skipping of exon 20 and to a corresponding reduction of ELP1 protein predominantly in the central and peripheral nervous system (CNS and PNS)(1–3). *ELP1* is one of the scaffolding subunits of the Elongator complex (4–6), a highly conserved protein complex involved in transcriptional elongation, tRNA modification and cytoskeletal remodeling (6–9). Several studies have demonstrated the role of ELP1 in neurogenesis, neuronal survival and differentiation as well as peripheral tissue innervation (10–20).

The major clinical features of FD are all due to progressive depletion of sensory and autonomic neurons (2, 21, 22). Common symptoms include gastrointestinal dysfunction, gastroesophageal reflux, vomiting crises, recurrent pneumonia, seizures, gait abnormalities, kyphoscoliosis, postural hypotension, hypertension crises, absence of fungiform papillae on the tongue, decreased deep-tendon reflexes, defective lacrimation and impaired pain and temperature perception (3, 23–29) Despite this complex neurological phenotype, FD patients also suffer from progressive visual dysfunction that severely affects their quality of life (30–32). Initially, it was reported that the loss of vision in FD patients resulted from corneal opacities, neovascularization and sensory defects such as corneal analgesia, severe dry eye, ulceration healing and incomplete closure of eye lids (33–37). However, recent detailed studies have shown that decreased visual acuity, loss of central vision, and temporal optic nerve pallor occur in FD patients even without any corneal complications, suggesting a neuro-ophthalmic nature of the disease (31). In FD, visual impairment is usually early onset and often progresses to legal blindness in the third decade of life (30). Individuals with FD show a significant reduction in the thickness of the retinal nerve fiber layer (RNFL) due to death of retinal ganglion cells (RGCs) (30–32). FD shares several similarities with other optic neuropathies caused by mitochondrial gene mutations such as Leber’s hereditary optic neuropathy (LHON) and dominant optic atrophy (DOA) (31).

Many efforts have been undertaken in the field to develop therapies aimed at correction of *ELP1* splicing defects, including splicing modulator compounds (SMCs), antisense oligonucleotide (ASO) and modified exon-specific U1 small nuclear RNA (38–40). Previously, our team has identified the small molecule kinetin (6-furfurylaminopurine) to be an orally active splicing modulator of *ELP1* both *in vitro* and *in vivo* (41, 42). As part of the NIH Blueprint Neurotherapeutics Network we have created a new class of highly potent SMCs, using kinetin as a starting molecule, and identified BPN-15477, a more potent and efficacious *ELP1* splicing modulator (43, 44). Despite this incredible progress, the field has not yet developed a therapy to prevent retinal degeneration in FD. The creation and characterization of mouse model able to recapitulate the retinal phenotype is the first step toward this effort. Two different *Elp1* conditional knock-out (CKO) mice with retinal degeneration were previously described, the *Tuba1a-cre; Elp1^flox/flox^* and the *Pax6-cre; Elp1^flox/flox^* mice (45, 46). In the *Tuba1a-cre; Elp1^flox/flox^* mouse, *Elp1* is deleted in the nervous system and in the *Pax6-cre;Elp1^flox/flox^* mouse, *Elp1* is deleted specifically in the retina (45, 46). Both these mice exhibit loss of RGCs and they are useful models to investigate the retinal pathology associated with complete loss of *Elp1*. However, since FD is caused by a reduction, not loss, of ELP1, they do not precisely model the human molecular defect and therefore they cannot be used to evaluate the *in vivo* efficacy of splicing modulator therapies. Most recently, we have generated a new phenotypic mouse model for FD by introducing the human *ELP1* transgene with the FD splice mutation (*TgFD9*) into a *Elp1* hypomorphic mouse (*Elp1^∆20/flox^*) (47, 48). The TgFD9; *Elp1^∆20/flox^* mouse recapitulates the same tissue-specific mis-splicing observed in FD patients and the major neurological symptoms of the disease (49). In the present study, we have characterized the progression of retinal disease in this FD mouse model and identified quantifiable retinal phenotypes which can be used to evaluate the efficacy of splicing targeted therapeutics in preventing retinal degeneration in FD.

## Results

### The FD phenotypic mouse recapitulates the optic neuropathy observed in FD patients

The RNFL is composed of non-myelinated axons of RGCs, which converge into the optic nerve. In several hereditary optic neuropathies, including glaucoma, the thinning of RNFL thickness results from damage of RGC axons which precedes the loss of RGCs (50–57). Many studies have demonstrated that patients with FD exhibit reduction in thickness of the RNFL layer due to the death of RGCs (30–32), and this loss is more profound near the temporal region of the optic nerve, specifically in the maculo-papillary region. Recently, we have generated a novel FD phenotypic mouse model, the *TgFD9; Elp1^∆20/flox^* mouse, by introducing a transgene carrying the human *ELP1* gene with the FD splice mutation (*TgFD9*) into an hypomorphic mouse that expresses low levels of endogenous *Elp1* (*Elp1^∆20/flox^*) (47–49). For simplicity, we will refer to TgFD9; *Elp1^∆20/flox^* mouse as the FD phenotypic mouse throughout the manuscript. Seeking to develop therapies targeting retinal degeneration in FD, we comprehensively characterized the progression of the retinal phenotype in this FD mouse model. We measured the thickness of retinal nerve fiber layer (RNFL) and ganglion cell-Inner plexiform layer (GCIPL) in the superior, inferior, nasal and temporal hemispheres of the retina using high-definition spectral domain optical coherence tomography (SD-OCT) (Figure 1A). At 3 months of age, we observed a significant reduction of RNFL and GCIPL layers in the temporal portion of the FD retina, as previously observed in FD patients. Starting from 6 months of age the thickness of RNFL and GCIPL layer becomes more uniformly reduced throughout the retina (Figure 1B and C). The thinning of these layers further progresses at 12 and 18 months of age in FD mice compared to control mice (Figure 1B and C). Even though, reduction of RNFL and GCIPL has been initiated in the temporal region of the retina, we observed uniform thinning of these layers throughout the retina after 3 months of age. These results indicate that our mouse model recapitulates the reduction of retinal thickness observed in FD patients.

**Figure 1:**
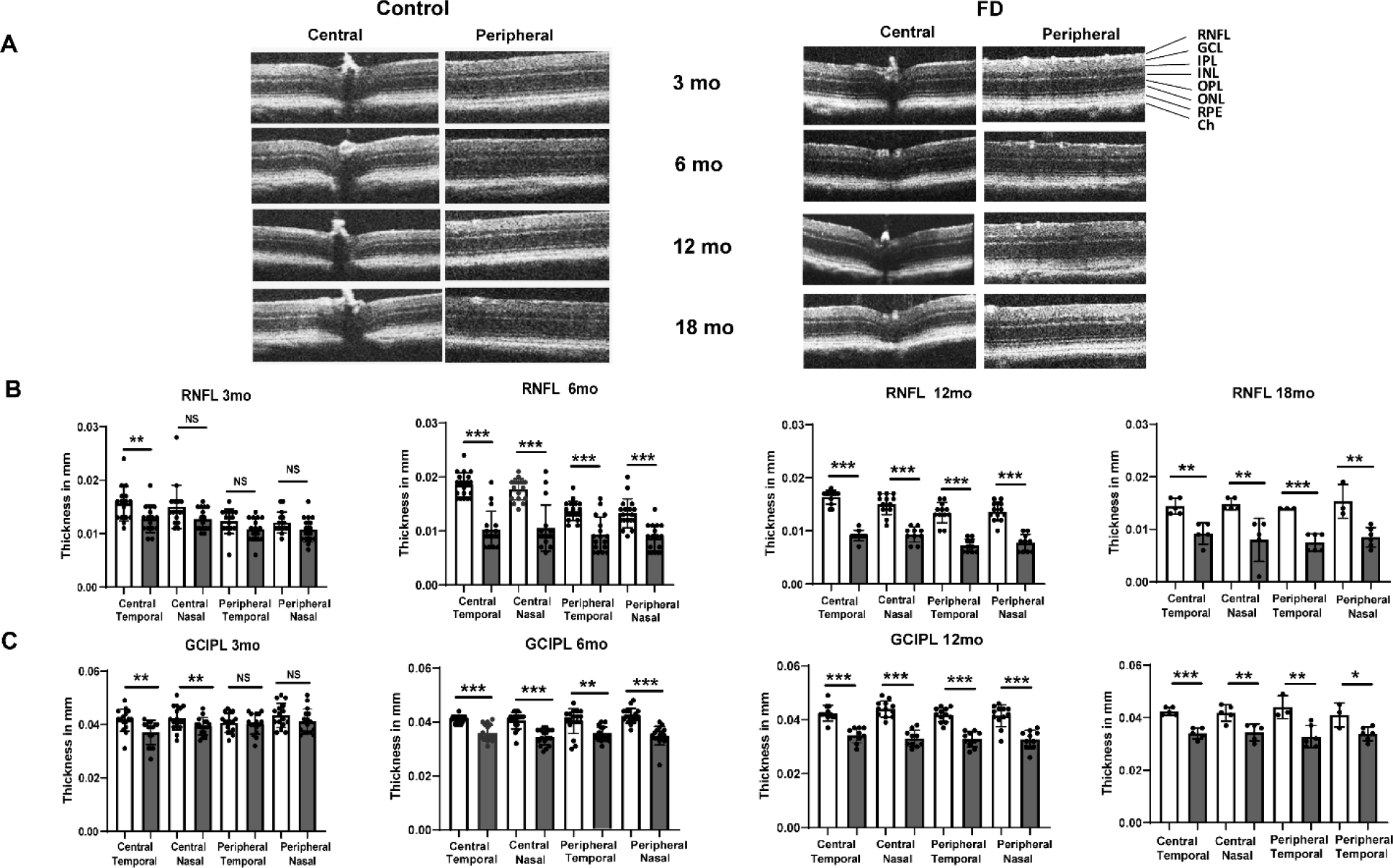
Measurement of RNFL and GCIPL thickness in FD mice using SD-OCT. **A.** Representative SD-OCT b-scan images of FD and control retinae at 3, 6, 12 and 18 months. **B.** Quantification of thickness measurements of RNFL in both central and peripheral regions of the retina in nasal and temporal hemispheres at 3 months (n=8), 6 months (n=9), 12 months (n=6), and 18 months (n=3) of age. **C.** Measurement of GCIPL thickness in both central and peripheral regions of the retina in nasal and temporal hemispheres at 3 months (n=8), 6 months (n=9), 12 months (n=6), and 18 months (n=3) of age. *P< 0.05, **P< 0.01 and ***P< 0.001. Error bars represent SEM. *P* value was calculated using students *t*-test.

### Selective loss of RGCs in the FD mouse retina

To evaluate if the reduction of RNFL and GCIPL thickness in the FD mice was due to loss of RGCs, we performed RGC counts in superior, inferior, nasal and temporal regions of the retina using retinal whole-mount analysis. We stained retinae from 3, 6- and 18-month-old mice using the RGC specific marker RNA-binding protein with multiple splicing (RPBMS) and we counted the number of RPBMS^+^ cells sited at 1mm from the optic nerve head (ONH) (Figure 2A and 2B). At 3 months of age, although we observed a trend toward reduction in RPBMS^+^ cells in the FD mice, the difference when compared with the control mice was not significant (Figure 2C). The RGC loss further progresses and, at 6 months, the number of RPBMS^+^ cells became significantly lower in the FD mice in the temporal (*P* < 0.002), nasal (*P* < 0.001), superior (*P* < 0.005) and inferior (*P* < 0.006) regions of the retina. At 12 months of age, the number of RPBMS^+^ cells in the FD retinae, although significantly lower than in the control mice, is comparable with the 6-month-old counts, indicating that the major RGC loss occurs between 3 and 6 months of age (Figure 2C). As expected, the control mice showed an age-dependent decline in the number of RGCs at 18 months. These results confirmed that the reduction in RNFL and GCIPL thickness results from progressive loss of RGCs.

**Figure 2:**
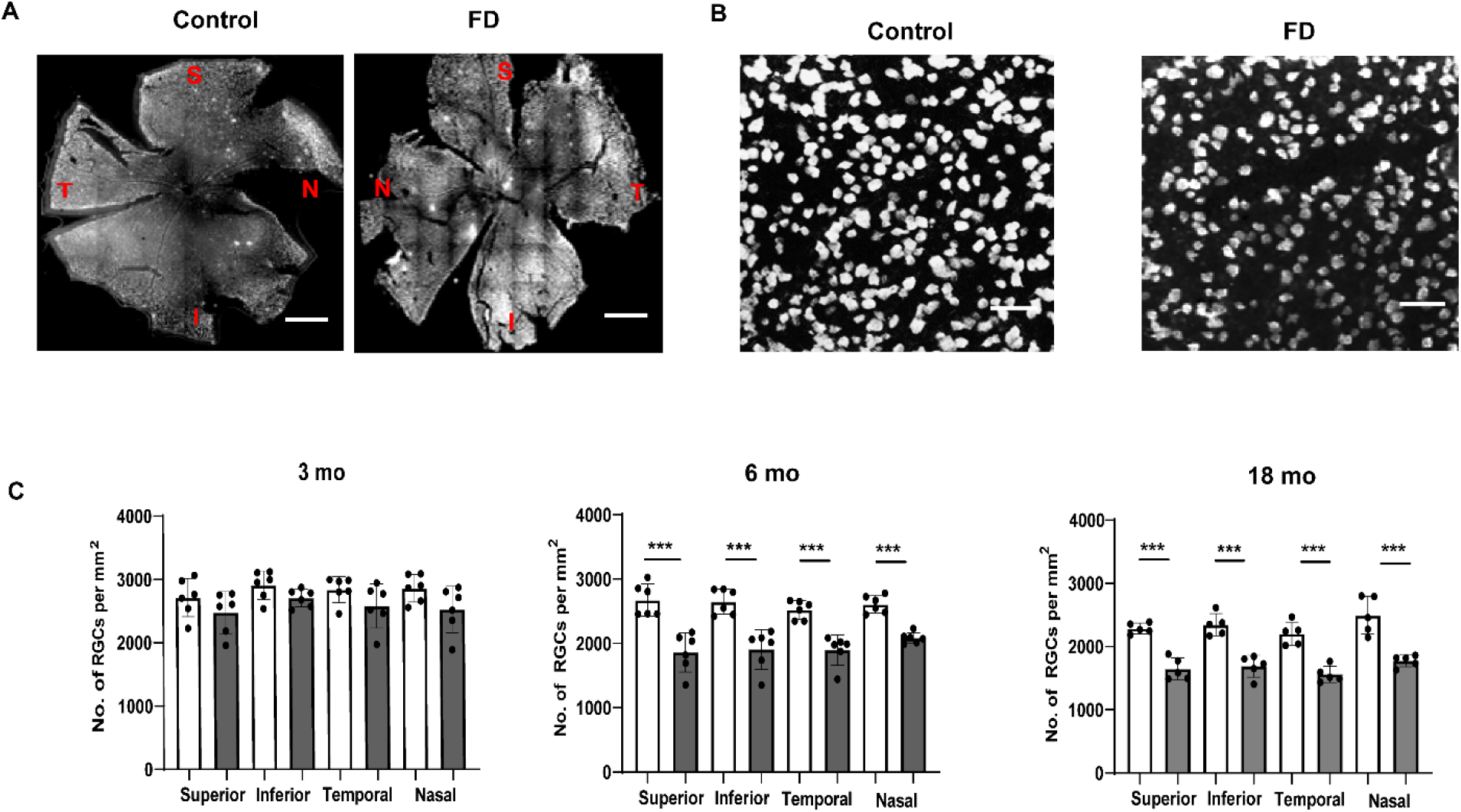
Progressive degeneration of RGCs in the phenotypic FD mouse model. Representative retinal whole-mount images from control and FD mice stained with RGC marker RBPMS. Scale bars, 100μm. RPBMS+ cells in control and FD retinae were counted in each quadrant in superior (S), inferior (I), nasal (N) and temporal regions at 1mm from the optic nerve head (ONH) at 3, 6 and 18 months of age. **B.** Higher magnification representative RPBMS staining images of control and FD retinae. Scale bars, 100μm. **C.** Bar plots of RPBMS^+^ cell counts in 3 month- (n=5), 6 month- (n=6) and 18-month-old (n=6) retinae from control and FD mice. Significant reduction in the number of RGCs was observed starting at 6 months of age and progresses at 18 months in FD mice compared to control mice. *P< 0.05, **P< 0.01 and ***P< 0.001. Error bars represent SEM. *P* value was calculated using students *t*-test.

Patients with FD do not show any significant photoreceptor loss; however, degeneration of photoreceptors was observed in *Tuba1a-cre; Elp1^flox/flox^* mice. To investigate whether RGC loss led to photoreceptor degeneration, we analyzed the number of photoreceptors in our FD mice and we did not observe significant changes at any age points tested (Supplementary figure 1).

### Optic nerve degeneration in the FD phenotypic mice

Histopathological analysis of the optic nerve from FD patients indicates diffuse degeneration of axonic bundles (30). To investigate if our FD mouse model also recapitulates this aspect of the disease, we collected optic nerves and retinae from FD mice and stained the axonal structures and the cell bodies of RGCs using the neurofilament (NF) antibody SMI-32. We performed this analysis at 8 months of age because our previous results have shown that the major RGC loss in FD mice occurs within the first 6 months of age. NF-staining of longitudinal optic nerve sections revealed that while the optic nerve fibers in the control mice were long, parallel and undistorted, the fibers in the FD optic nerve had significantly more retraction bulbs and holes (Figure 3A). We therefore morphologically evaluated these alterations in the optic nerve sections using the SMI-32 score (58). This score assesses the structural integrity of the optic nerve fibers. It is usually calculated based on SMI-32 staining and it is graded as 0 for intact structure and no retraction bulbs, 1 for short axons and occasional retraction bulbs and 2 for numerous retraction bulbs and holes. The SMI-32 score of the FD optic nerves was 1.5, two times the score of the control optic nerve (*P* < 0.032; Figure 3B). We then investigated whether these alterations led to thinning of the optic nerve by measuring the circumference of the DAPI-stained optic nerve cross sections (Figure 3C). We found that the circumference of FD optic nerves was significantly smaller compared to the control optic nerves (*P* < 0.025; Figure 3C and D). Finally, in order to analyze the number of axonal bundles entering the optic nerve, we performed NF staining of whole-mount retinae (Figure 3E and F). We counted the axonal nerve bundles entering the optic nerve in superior, inferior, nasal and temporal hemispheres. Interestingly, the number of axonal bundles in the FD retinae was significantly lower compared to the control retinae in every retinal region tested (Superior: *P* < 0.007, inferior:0.0001, temporal:0.0005, nasal:0.005; Figure 3G). Overall, our findings demonstrate that the FD mouse correctly models the optic nerve degeneration observed in FD patients.

**Figure 3:**
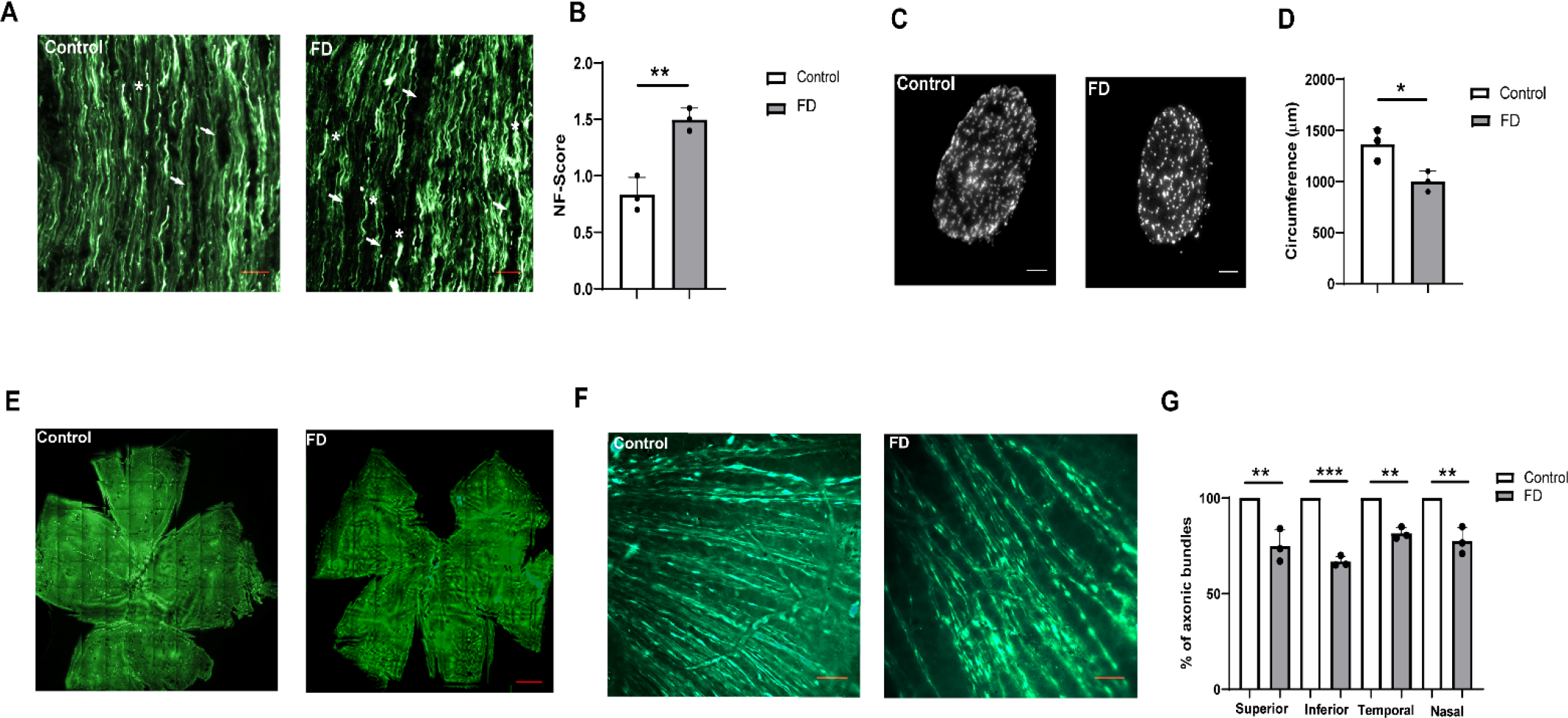
Degeneration of the optic nerve was observed in the FD phenotypic mouse. **A.** Representative optic nerve sections in 8-month-old FD and control retinae stained with NF (SMI-32) antibody. NF staining indicate numerous retraction bulbs, shorter axons and holes in the optic nerve tissue of FD mice when compared with control mice. Scale bars, 100μm. Arrows indicate holes and *indicate retraction bulbs in the image. **B.** Graph representing the scoring of SMI-32-stained optic nerve sections revealed significant degeneration of optic nerve evidenced by structural distortion of the optic nerve (*p<*0.032: *n* =3). **C.** Staining of optic nerve cross sections at 8 months using DAPI. Scale bars, 50μm. **D.** Bar plot representing the measurement of the optic nerve circumference using image J. Significant reduction in the circumference of the optic nerve was observed in FD mice at 8 months (*P<* 0.025: n=3). **E.** Representative picture of NF (SMI-32) staining of retinal whole-mounts from mice at 8 months of age. Scale bars, 50μm. **F.** Higher magnification image indicates axonic bundles surrounding the optic nerve. Some RGCs were also observed with NF staining. **G.** Measurement of number of axonic bundles surrounding the ONH. Axonic bundles were counted 0.5 mm from the ONH in 1mm square at superior (*P* < 0.007), inferior (*P* <0.0001), nasal (*P*<0.0005) and temporal (*P*<0.005) regions (n=3).

### *ELP1* exon 20 inclusion is lower in RGCs compared to other retinal cells

Although *ELP1* is ubiquitously expressed, the major FD splicing mutation leads to a tissue-specific skipping of *ELP1* exon 20 with the lowest amount of full-length *ELP1* expression in the CNS and PNS (20, 47, 49, 59). In order to assess the splicing of the human *ELP1* transgene in different regions of the eye, we collected lens, cornea, retina, iris, optic nerve and posterior eye cup from the transgenic mice *TgFD9* (47). For this analysis, we specifically used the *TgFD9* mice because it is a suitable model to assess the FD splicing defect *in vivo as* the splicing pattern of the human *ELP1* transgene is identical between the *TgFD9* and the *TgFD9; Ikbkap*^∆20/flox^ mice (47, 49) (Supplementary figure 4). As shown in Figure 4, the amount of exon 20 inclusion was lower in the cornea and optic nerve when compared to other ocular tissues tested (Figure 4A and 4B). Surprisingly, although the retina is part of the CNS, it was not the ocular tissues with the lowest amount of full-length *ELP1* transcript expression. We therefore hypothesized that the mutant *ELP1* transcript might splice differently in distinctive retinal cells and assessed *ELP1* splicing in RGCs, because they are the most affected cell type in the FD (Figure 4C). As RGCs comprise only 1% of the total retinal cells, the *ELP1* splicing analysis of the entire retina might not necessarily represents the splicing efficacy of this specific subpopulation of neurons. RGCs were enriched from *TgFD9* retinae with fluorescence activated cell sorting (FACS) using positive selection for the anti-thymocyte antigen (Thy1/CD90.2). Interestingly, exon 20 inclusion was significantly reduced in RGCs when compared to other retinal cells (Figure 4C, D and supplementary figure 3) supporting the idea that the RGC loss observed in the FD retinae might result from ELP1 amounts falling below cell-specific threshold.

**Figure 4:**
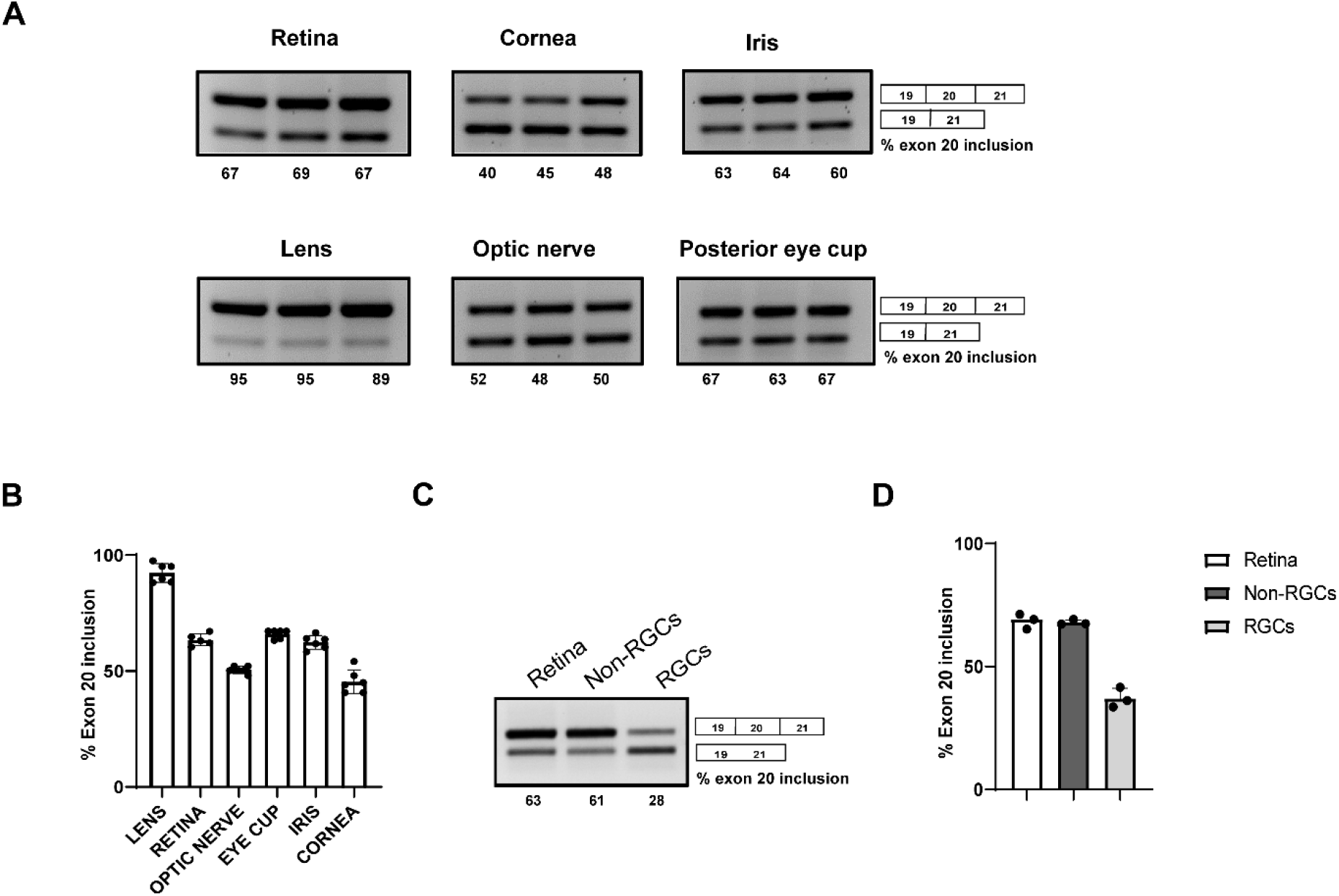
Ocular- and cell-specific mis-splicing of the human FD *ELP1* transgene. **A.** Splicing analysis of human *ELP1* transcripts in the retina, lens, optic nerve, iris, cornea and posterior eye cup of 3 to 4-month-old FD mice. **B.** Quantification of the percentage of exon 20 inclusion in various ocular tissues (n=6). **C.** Splicing analysis of human *ELP1* transgene in RGCs and non-RGC retinal cell population compared to whole retinal tissue. **D.** Quantification of percentage of exon 20 inclusion in RGCs compared to other retinal cells (n=3).

### The small-molecule BPN-15477 corrects*ELP1* splicing defect in the mouse retina

Our work has been focused on the development of small molecule SMCs to correct *ELP1* splicing defect in FD. We have shown that daily consumption of kinetin (6-furfuryl amino purine) rescues neuronal phenotype in our FD mouse model and, more recently, we have identified a novel SMC, BPN-15477, significantly more potent and efficacious than kinetin (40, 43). However, we have never tested the ability of our SMCs to correct *ELP1* splicing in the retina. Prior to initiating preclinical trials in our phenotypical FD mouse model, we sought to determine if BPN-15477 could modulate exon 20 inclusion in the retina of the *TgFD9* mouse. This mouse, although phenotypically normal, perfectly recapitulates the *ELP1* splicing defect observed in patients and therefore is the ideal model to test the effect of SMCs on splicing *in vivo* (49). BPN-15477 was administered to the *TgFD9* mice starting at birth through special formulated chow. The special chow was formulated so that each mouse received 70 mg/kg/day. At birth, pups were randomly assigned to vehicle- or BPN-15477-treated groups and were maintained in the same treatment regime till time of sacrifice. To assess *ELP1* mRNA splicing, RT-PCR assays were performed on total RNA extracted from brain and retina of the mice carrying the *TgFD9* transgene. Remarkably, BPN-15477 treatment significantly improved *ELP1* exon 20 inclusion in the brain and fully corrected *ELP1* splicing in the retina in the BPN-15477-treated mice when compared to the vehicle-treated mice (Figure 5). This result demonstrated that our splicing modulator BPN-15477 efficiently crosses the blood retinal barrier (BRB) and therefore can be used to correct *ELP1* splicing defect in the retina.

**Figure 5:**
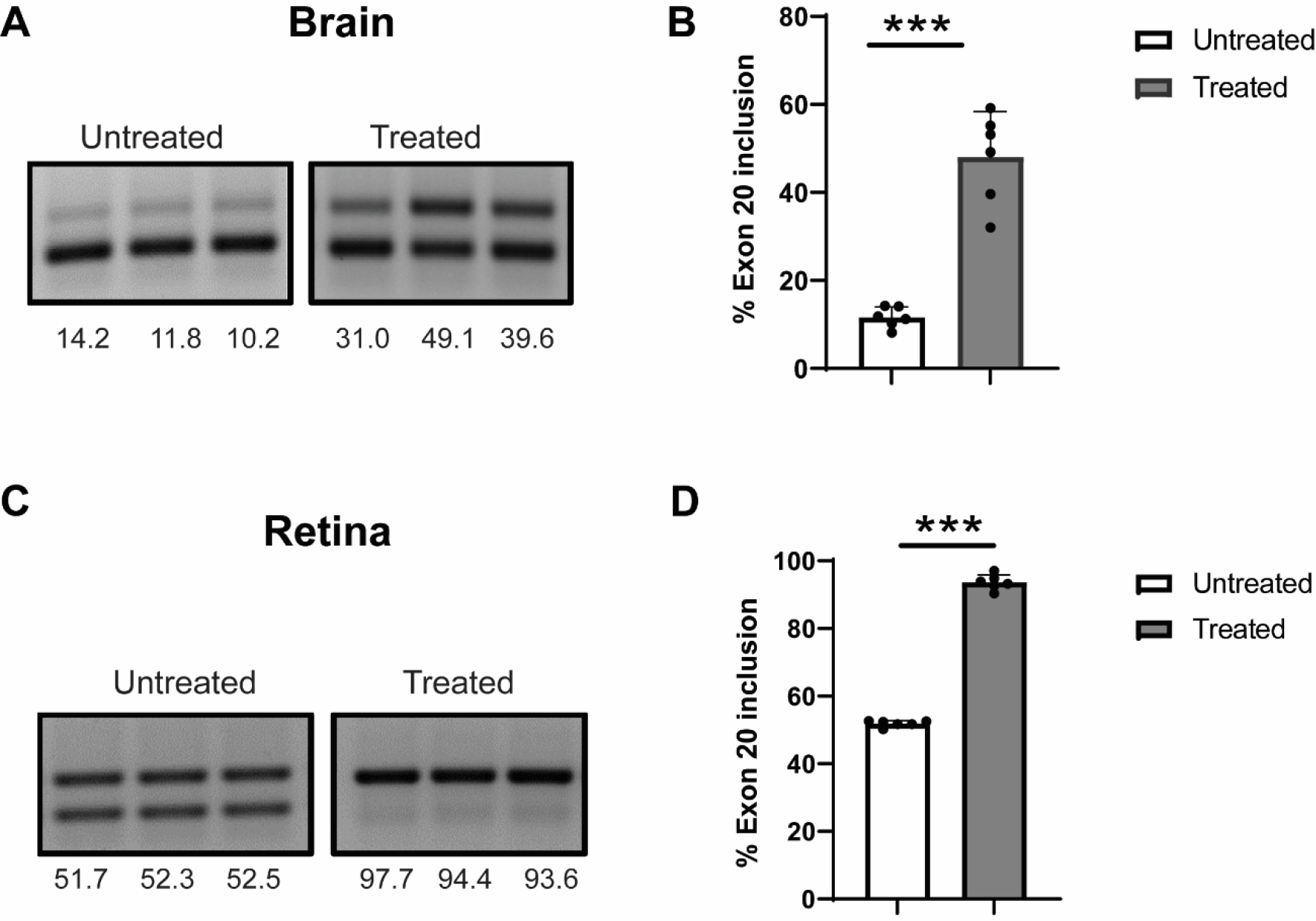
BPN-15477 treatment improves exon 20 inclusion in the retina and brain of mice carrying the human FD transgene *TgFD9*. **A.** Representative gel picture indicating splicing analysis of human *ELP1* transcript in the brain from *TgFD9* mice treated with −70 mg/Kg/day of BPN-15477. **B**. Quantification of exon 20 inclusion in the brain (n=6; *P*<0.0001). **C.** Representative gel picture of splicing analysis of human *ELP1* transcript in the retina. Note the near complete correction of *ELP1* splicing in *TgFD9* mice after BPN-15477 treatment **D.** Quantification of exon 20 inclusion in the retina (n=6; *P*<0.0001).

## Discussion

FD is a devastating neurodegenerative disorder caused by a splicing mutation in the *ELP1* gene. Despite the severe sensory and neurological manifestations, one of the most debilitating aspect of the disease is progressive visual loss. Previous studies using high-definition OCT showed that all FD patients suffer from an optic neuropathy featured by reductions of the RNFL which is due to loss of the macular RGCs (30, 60). Although, pharmacological and surgical supportive treatments are available to alleviate some of the disease symptoms (61), to date there are no therapeutic interventions aimed at stopping vision loss and most of the patients become visually impaired or legally blind after their third decade of life. We have comprehensively characterized the retinal pathology of the FD phenotypic mouse model using a combination of *in vivo* imaging and histological analyses and show that this mouse accurately models the human phenotype. Using SD-OCT, we have shown that the reduction of RNFL and GCIPL layers in the FD mice starts as early as 3 months of age and progresses further at 6 months, recapitulating the retinal disease progression observed in FD patients (30). Consistent with the degeneration of RNFL and GCIPL layers, the FD mouse also showed progressive retinal ganglion cell loss after 3 months of age, confirming that ELP1 is not required for RGC development and offering a window for therapeutic intervention (46). Although *ELP1* is expressed in all major retinal neurons (45), we have shown that the retinal degeneration in FD is specific to RGCs and no significant differences in the photoreceptor number or in glial activation were observed even in old mice. Moreover, we have proven that our phenotypic mouse correctly models the optic nerve degeneration observed in FD patients. This extensive characterization allowed us to identify quantifiable endpoints that can be used for future pre-clinical trials aimed at testing the efficacy of existing or novel disease modifying therapies targeting the retina.

Many efforts have been undertaken in the field to better understand the pathophysiological mechanisms underlying retinal degeneration and to develop disease modifying therapies for FD. The majority of these approaches aim to correct the *ELP1* splicing defect and include SMCs, ASO and modified exon-specific U1 small nuclear RNA (38–40). Gene replacement strategies have been hindered by the large size of the *ELP1* gene. The creation and characterization of a mouse model able to recapitulate the retinal phenotype now provides a crucial resource for testing all of these potential therapies. The previously described *Elp1* CKO mice, *Tuba1a-cre; Elp1^flox/flox^* and *Pax6-cre; Elp1^flox/flox^*, are useful models to investigate the retinal pathology associated with complete loss of *Elp1*, however, they do not accurately model the molecular cause of the disease and therefore they cannot be used to evaluate the *in vivo* efficacy of splicing modulator therapies (45, 46). Conversely, the retinal phenotype of the *TgFD9; Elp1*^∆20/flox^ mouse recapitulates the pathogenic mechanism of disease and therefore is the ideal model system to assess the *in vivo* efficacy of splicing correction in the retina. (49).

Splicing analysis of the human FD *ELP1* transgene in different eye regions showed that the highest amount of exon 20 inclusion was observed in the lens, while cornea and optic nerve were among the ocular tissues with lowest amount of full-length *ELP1* transcript. During embryonic development, lens and cornea are derived from the surface ectoderm, whereas retina and optic nerve extends from the diencephalon (62). We have previously shown that the amount of full length *ELP1* expression is significantly lower in the CNS and PNS when compared to other tissues both in FD patients and in mouse models (20, 47, 49). Therefore, we were surprised to find that the retina, although part of the CNS and largely affected by the disease, expressed 63% of *ELP1* full-length transcript. We then hypothesized that the mutant *ELP1* transcript could splice differently in different retinal cells and assessed *ELP1* splicing in RGCs because they are the most affected retinal neuronal subtype in FD. As anticipated, we found that *ELP1* exon 20 inclusion was significantly lower in this specific subpopulation of neurons when compared to the rest of the retina, demonstrating that the RGC loss in FD might result from *ELP1* amounts falling below a cell-specific threshold. Finally, in order to test the ability of our newly identified SMC to correct *ELP1* splicing in the retina, we administered BPN-15477 to the *TgFD9* mice starting at birth through special formulated chow (43). BPN-15477 fully corrects *ELP1* splicing defect in the retina, proving for the first time the therapeutic potential of an oral treatment to prevent retinal degeneration in FD. Taken together, the identification of quantifiable retinal phenotypes as well as the demonstration that BPN-15477 can correct *ELP1* splicing in the retina sets the stage for future pre-clinical testing that will move targeted treatments for blindness in FD to the clinic.

## Materials and Methods

### Animal maintenance

Mice used in this study were maintained according to the Association for Research in Vison and Ophthalmology (ARVO) statement for the use of animals in ophthalmic and vision research and with protocols approved by Institutional Animal Care and Use Committee of the Massachusetts General Hospital. The mice used for this study were housed in the animal facility of Massachusetts General Hospital (Boston, MA, USA), provided with constant access to a standard diet of food and water, and maintained on a 12-h light/dark cycle. For BPN-15477 treatment, assigned mice were fed on BPN-15477 containing chow. Mice were genotyped by PCR amplification of the genomic DNA obtained from tail biopsies and using the following primers: *Elp1* 1F (5′-TGATTGACACAGACTCTGGCCA-3′) and *Elp1* 4R (5′-CTTTCACTCTGAAATTACAGGAAG-3′) to discriminate the *Elp1* alleles and Tg Probe1 F (5′-GCCATTGTACTGTTTGCGACT-3′) and TgProbe1R (5′TGAGTGTCACGATTCTTTCTGC-3′) to discriminate the *TgFD9* transgene

### Spectral Domain Optical coherence tomography (SD-OCT)

For *in vivo* imaging of the retina, mice were anesthetized by placing them in a mobile isoflurane induction chamber and the vaporizer was set to an isoflurane concentration of 2% at 2 L/min O_2_. Pupils of the mice were dilated using 2.5% phenylephrine, 1% tropicamide. 0.5% proparacaine is used as a topical anesthetic during the procedure. SD-OCT imaging was performed using a Leica EnvisuR2210 OCT machine. Measurements were made 100μm from the optic nerve for central retina and the peripheral retina. Control and FD mice at 3, 6, 12 and 18 months of age were analyzed. Linear B-scans of superior, inferior, nasal and temporal retina were performed, and the thickness of the retina was measured using Bioptigen InVivoView Clinc software.

### RT-PCR analysis of*ELP1* transcripts

Following euthanasia of the mice, eyes were enucleated along with the optic nerve and placed in phosphate buffered saline (PBS) on ice. Dissection was performed to separate cornea, lens, iris, optic nerve, retina, and posterior eye cup from each eye and snap frozen using liquid nitrogen. The tissues were homogenized in ice-cold QIAzol (Qiagen) lysis reagent using a Tissue Lyser (Qiagen). Total RNA was extracted using the standard chloroform extraction procedure. Concentration of the total RNA for each sample was determined with a Nanodrop ND-1000 spectrophotometer. cDNA was synthesized by reverse transcription with approximately 0.5 μg of total RNA, Random Primers (Promega), and Superscript III reverse transcriptase (Invitrogen) according to the manufacturer’s instructions. To perform RT-PCR and detect *ELP1* exon 20 inclusion, 100 ng of starting RNA equivalent cDNA was subjected to PCR reaction with the GoTaq green master mix (Promega) and 30 amplification cycles (94°C for 30 s, 58°C for 30 s, and 72°C for 30 s). The human-specific forward and reverse primers spanning exon 19 to 21 of the *ELP1* gene 5′-CCTGAGCAG CAATCATGTG−3′ and 5′-TACATGGTCTTCGTGACATC-3′ were used for amplification both wild type and mutant isoforms of human *ELP1* transcript. The PCR products were resolved on 1.5% agarose gels containing ethidium bromide for detection. The relative amounts of normal and exon 20 excluded *ELP1* isoforms were quantified using ImageJ software, and the integrated density value for each band was determined and expressed as percentage as described previously(49).

### RGC Enrichment

RGCs were enriched from total retinal cells from a humanized transgenic mouse *TgFD9*. We followed previously published protocol with slight modifications for RGC enrichment(63). Briefly, a total of 10, 2-3-month-old *TgFD9* mice were enucleated, and the retinae were pooled for fluorescence activated cell sorting (FACS) analysis. Retinal cell suspension was prepared in a solution containing PBS with 1% FBS by gentle maceration and filtration by passing through a 70um nylon strainer. The retinal cells were immunolabelled using CD90.2AF-700, (1μg) CD48 PE-Cyanine 7(0.4μg), CD15 PE (0.02μg) and untagged CD57 (0.4μg). After 30 min incubation and washing cell suspension was incubated with BV421 (0.1μg) secondary antibody. After washing twice, labelled retinal cell suspension was subjected to FACS sorting to enrich CD90.2^+^, CD48^−^, CD15^−^, CD57^−^ phenotype.

### Immunohistochemistry

Eyes were enucleated and fixed in 4% paraformaldehyde overnight at 4°C. After a single PBS wash, eyes were cryoprotected in 30% sucrose overnight at 4°C and embedded in optimal cutting temperature compound (Sakura Finetek, Torrance, CA) and cryo-sectioned into 15μm thin sections. Sections were fixed in dry ice-cold acetone for 15 mins and washed with PBS 3 times. Antigen retrieval was then performed by incubating the sections using the buffer containing 0.1mtris.HCL ph 8, 50mM EDTA ph 8.0 and 20μg/ml proteinase K for 10 min at room temperature. After washing 2 times in PBS, retinal sections were blocked with animal-free blocker (Vector Laboratories, Burlingame, CA) containing 0.5% Triton X-100 for 1 h at room temperature, then primary antibodies anti-GFAP (Glial fibrillary acidic protein) (NeuroMab, Davis, CA), anti-Sox9 (SRY-Box Transcription Factor 9) (EMD Millipore, Billerica, MA) were applied and incubated at 4°C overnight in a moist chamber. Sections were washed three times with PBS with 0.1% triton X 100 and incubated with secondary antibodies (Invitrogen; Jackson Immuno Research, West Grove, PA) for 1 h at room temperature. Sections were mounted with hard-set mounting media containing DAPI (Vector labs) and fluorescent microscopy was performed.

### Retinal whole mounting and RGC counting

Fixation of the eyes was performed at room temperature for 1 hour in 4% PFA and eyes were marked with a yellow tissue marking dye on the temporal surface. After fixation, retinae were isolated, with each temporal retina marked with a small cut. Nonspecific binding was blocked by incubating with animal-free blocker containing 0.5% Triton X-100 overnight at 4°C, and anti-RPBMS antibody was applied overnight at 4°C. Retinae were incubated with secondary antibodies for 1 hour at room temperature and mounted on slides. Images were acquired using a LeicaDMi8 epifluorescent microscope and MetaMorph 4.2 acquisition software (Molecular Devices, San Jose, CA). Whole scans of complete flat mount samples were attained at 20X magnification using scan stage and autofocus. With ImageJ software, the number of RPBMS+ cells were measured (blinded) at an area of 1×1 mm at 1 mm square from the ONH at superior, inferior, temporal, and nasal hemispheres(46).

### Histological examination

Following euthanasia of the mice with CO2, eyes were marked with a yellow tissue marking dye on the temporal surface. Eyes were then enucleated and fixed overnight in 4% and embedded in paraffin for sectioning. Approximately 5μm sections were cut through the center of the eye based on the position of the optic nerve and subjected to hematoxylin and eosin (H&E) staining. Images were acquired by light microscopy using MetaMorph 4.2 software. Whole retina images were digitally developed using scan stage function for automated image patching. Morphology of the retina of both FD and control mice (n=5 each group) was analyzed with ImageJ. The number of rows of photoreceptor nuclei in a single column in the outer nuclear layer (ONL) was counted as described previously (45). Nuclei counting was performed at intervals of 0.25 mm starting at the optic nerve head (ONH) and toward both the temporal and nasal retinal hemispheres. Each sample was imaged and measured (blinded) in triplicate.

### Oral administration of BPN-15477 using specially formulated diet

FD mice were fed special diet containing 0.08% of BPN-15477 (LabDiet 5P00 with 800 ppm BPN-15477), corresponding to a daily consumption of 70 mg/ Kg/ day from birth until 5 weeks of age. In the control group, *TgFD9* transgenic mice were fed with vehicle diet (LabDiet 5P00). The mice were given water ad libitum.

### Statistical analysis

The data were statistically analyzed using graph pad prism software (La jolla CA). Comparison between the two groups was performed using two-tailed independent Student’s t-tests. The data are presented as the mean – standard error. P values <0.05, <0.01, and <0.001 were considered as significant.

## Acknowledgements

We thank Dr. Horacio Kaufmann at the Dysautonomia Treatment and Evaluation Center at New York University Medical School, Dr. Frances Lefcort at Montana State University and Dr. Carlos Mendoza at the Bascom Palmer Eye Institute for their long-standing collaboration and helpful discussions. This work was supported by National Eye Institute (NEI) grant (1R01EY029544-01 to S.A.S. and L.H.V.) and by Grousbeck Family Foundation (to L.H.V).

## Conflict of Interest Statement

The authors declare competing financial interests.

Susan A. Slaugenhaupt is a paid consultant to PTC Therapeutics and is an inventor on several U.S. and foreign patents and patent applications assigned to the Massachusetts General Hospital, including U.S Patents 8,729,025 and 9,265,766, both entitled “Methods for altering mRNA splicing and treating familial dysautonomia by administering benzyladenine,” filed on August 31, 2012 and May 19, 2014 and related to use of kinetin; and U.S. Patent 10,675,475 entitled, “Compounds for improving mRNA splicing” filed on July 14, 2017 and related to use of BPN-15477. Elisabetta Morini and Susan A. Slaugenhaupt are inventors on an International Patent Application Number PCT/US2021/012103, assigned to Massachusetts General Hospital and entitled ,RNA Splicing Modulation- related to use of BPN-15477 in modulating splicing. Luk H. Vandenberghe (LHV) holds equity in Affinia Therapeutics and Akouos and serves on the Board of Directors of Affinia Therapeutics, Addgene and Odylia. LHV is compensated for his scientific advisory position with Affinia and Akouos. LHV is an SAB member to Akouos, consultant to Affinia and Novartis and receives research support from Novartis. LHV’s interests were reviewed and are managed by Mass Eye and Ear and Mass General Brigham in accordance with their conflict-of-interest policies.

## Abbreviations

FBS: Fetal bovine serum
DAPI: 4′,6-diamidino-2-phenylindole
HCL: Hydrochloric acid
EDTA: Ethylenediaminetetraacetic acid
mRNA: messenger Ribonucleic acid
Tg: Transgenic

