## Supplementary figures for "Selective retinal ganglion cell loss and optic neuropathy in a humanized mouse model of familial dysautonomia"

**Supplementary information**

**Supplementary Figure 1**

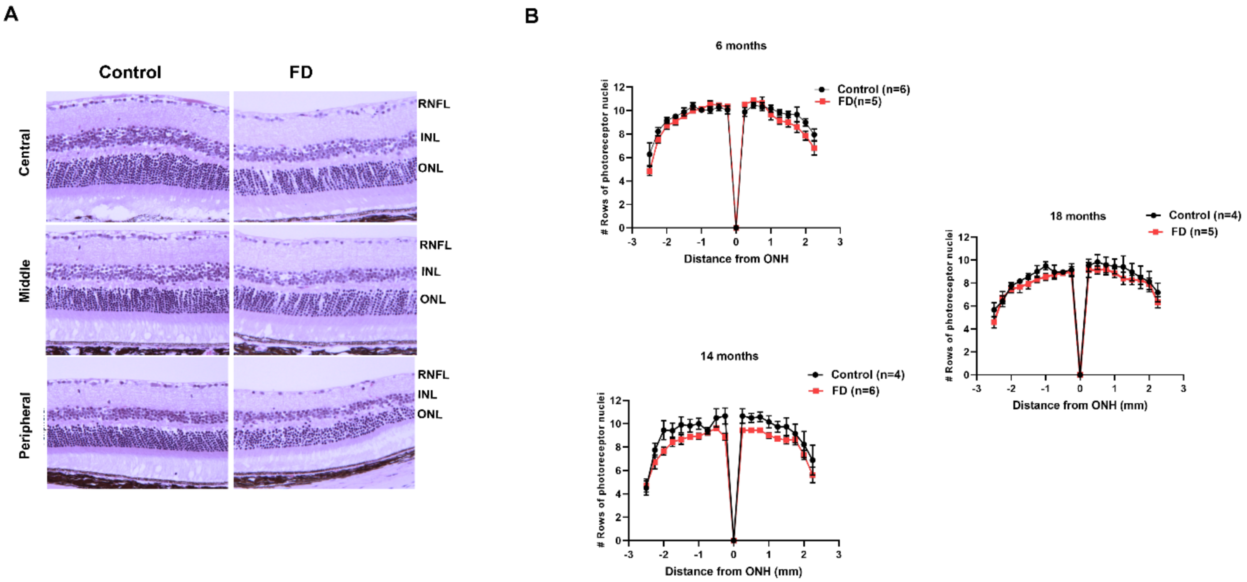


**Supplementary figure 1. No loss of photoreceptor was observed in the FD phenotypic mice:** Representative image of H&E stained retinal cross sections at central (0.25mm from ONH), middle (1mm from ONH) and peripheral (1.75mm from ONH). Overall structure of the retina has appeared to be grossly normal. Graphical representation of counts of the number of rows of photoreceptor nuclei in the outer nuclear layer. Nuclei counts were made 0.25mm from ONH in both temporal and nasal retinae at 6, 14 and 18 months of age in control and FD mice. No significant difference in photoreceptor numbers between control and FD retinae was observed. Statistical significance was analyzed by students t-test (n=at least 4 in control retinae and 5 for FD retinae).

**Supplementary figure 2**

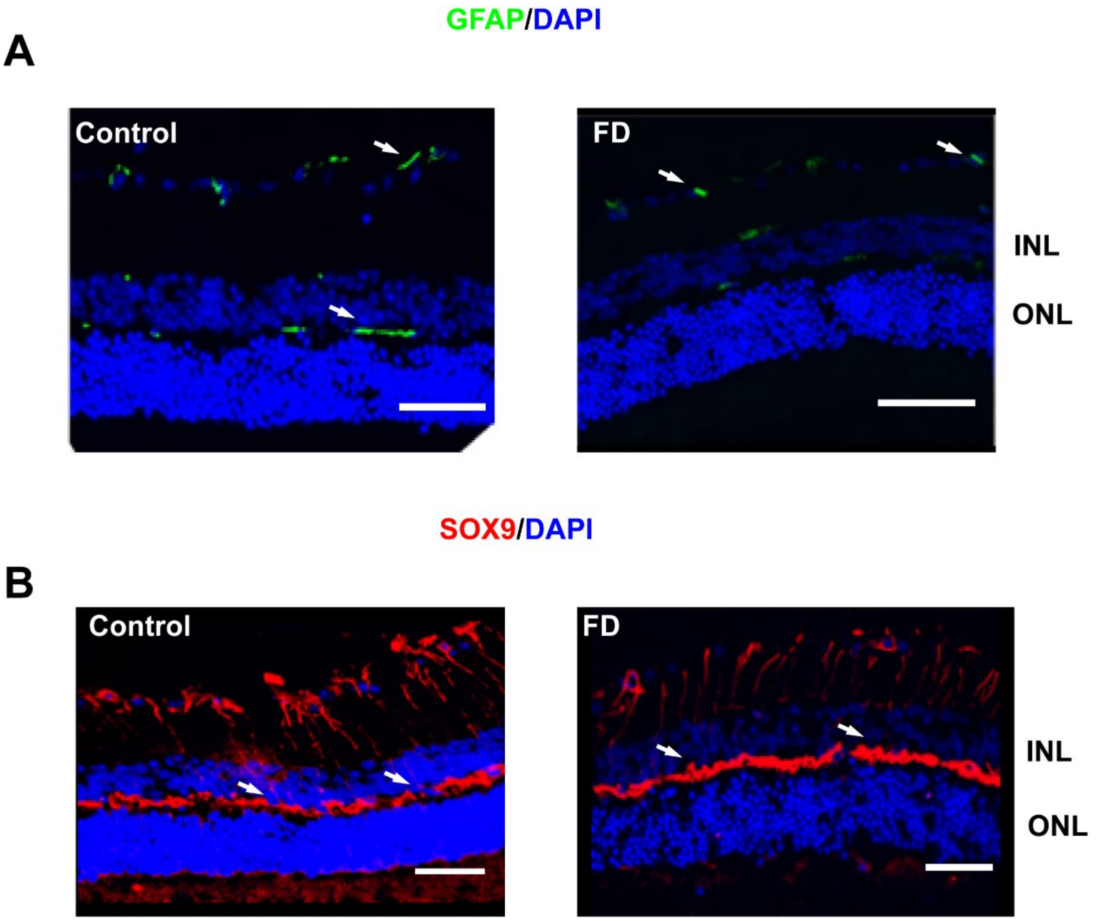


**Supplementary figure 2. No activation of glial cells was observed in FD phenotypic mice:** Immunohistochemical analysis of control and FD retinae. Representative images indicating staining of **A**. GFAP (green) (Scale bars, 500µm). and **B.** Sox-9 (Red) (Scale bars, 400µm). Nuclei were counter stained using DAPI (blue). No difference in glial activation was observed between FD and control retinae. Sox9 staining pattern also appeared normal indicating no differences in activated Mueller cells between control and FD retinae.

**Supplementary figure 3**

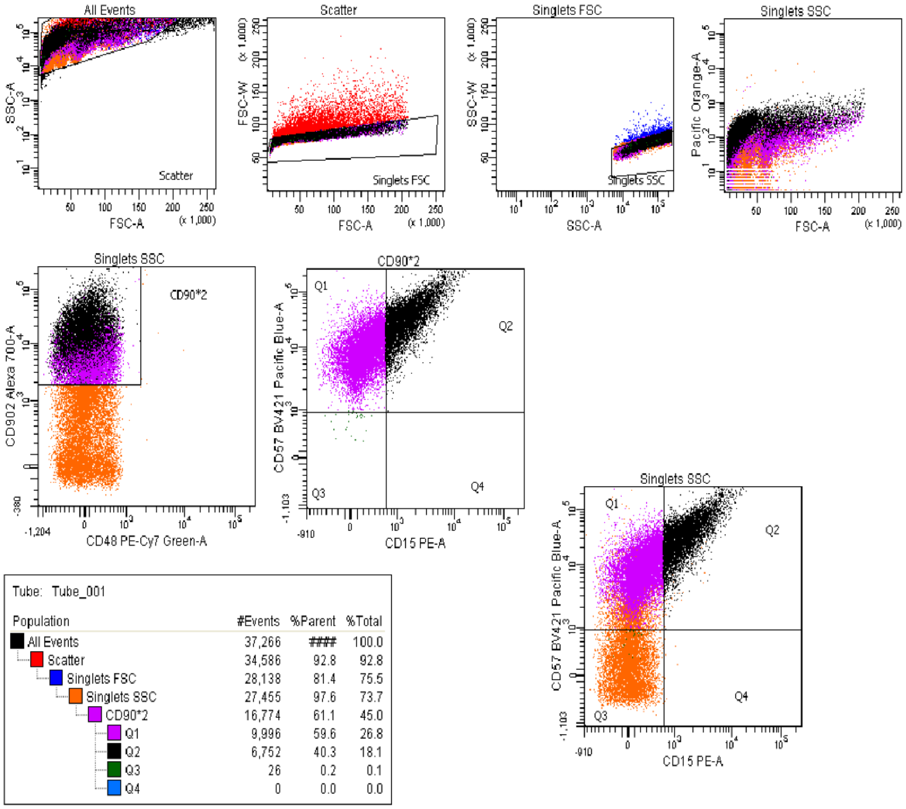


| Population | #Events | %Parent | %Total |
|---|---|---|---|
| All Events | 37,266 | #### | 100.0 |
| Scatter | 34,586 | 92.8 | 92.8 |
| Singlets FSC | 28,138 | 81.4 | 75.5 |
| Singlets SSC | 27,455 | 97.6 | 73.7 |
| CD90*2 | 16,774 | 61.1 | 45.0 |
| Q1 | 9,996 | 59.6 | 26.8 |
| Q2 | 6,752 | 40.3 | 18.1 |
| Q3 | 26 | 0.2 | 0.1 |
| Q4 | 0 | 0.0 | 0.0 |

**Supplementary figure 3. Isolation of RGCs from the total retinal population using FACS:** Sorting strategy was based on the inclusion of CD90.2 cells and exclusion of CD48, CD15 AND CD57 cells. Overview of cell population was obtained by Plot size indicated by FSC and internal complexity indicated by SSC. Initial gating was set up to discriminate single cells and aggregates using FSC-W versus FSC-A and SSC-W and SSC-A. Retinal cells were labelled with AF700 conjugated CD90.2, PE-Cyanine7- conjugated CD48, PE-conjugated CD15, and CD57 was probed using BV421 antibody. Next gating strategy was set up for the selection of CD90.2+ and CD48- cells. This is followed by CD57 versus CD15 plot was generated using the previously selected CD90.2+, CD48- population.

**Supplementary figure 4**

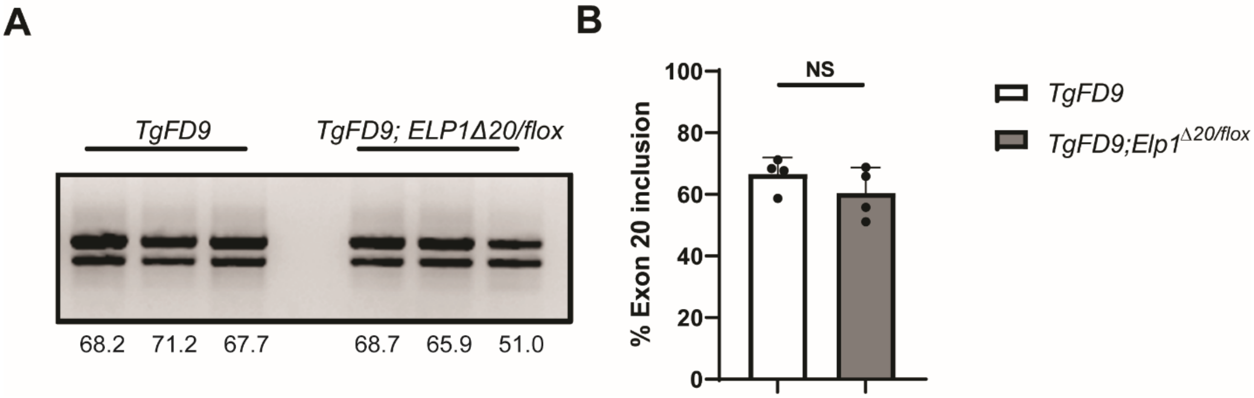


**Supplementary figure 4**. **Splicing analysis of the human FD *ELP1* transgene in *TgFD9* and *TgFD9; ELP1*$^{\Delta 20/flox}$ mice: A.** Representative gel picture indicating splicing analysis of human *ELP1* transcript. **B**. Quantification of exon 20 inclusion in *TgFD9* and *TgFD9; ELP1*$^{\Delta 20/flox}$ (*n=4*). No significant difference was observed in exon 20 inclusion between *TgFD9* and *TgFD9; ELP1*$^{\Delta 20/flox}$ mice.
